# Female assortment in Gouldian Finch (*Chloebia gouldiae*) Flocks: A case of female-driven social niche construction?

**DOI:** 10.1101/2025.06.22.660960

**Authors:** Gregory Kohn, Thomas Mateo, Alex Gibson, Christina Galloway

## Abstract

The organization of animal groups is both a cause of and a consequence of patterns of interactions among individuals. The stability of social organization reflects how individuals construct and maintain resilient patterns of interactions across changes in group size and membership. Who you choose to associate with may have numerous advantages, primarily through avoiding potentially costly interactions with others. In this study, we describe patterns of social interactions in multiple flocks of Gouldian finches that were exposed to changing social conditions. In the first condition, flocks experienced repeated introductions of juveniles into the group. In a second condition, an adult flock was exposed to changing conditions where males were swapped between the flocks. In almost all flocks, we found significant levels of female-driven sex assortment. Further analysis shows that females were more likely to “leave” interactions after being approached by males than males, thus constituting a social decision to avoid male interactions. These findings highlight how the stable interactive decisions made by individuals contribute to higher-order patterns of organization in bird flocks.

## 1 Introduction

Most social species inhabit loosely cohesive fission-fusion societies, where group size and composition fluctuate over short timescales. The social organization of these societies reflects the ability of individuals to reconstruct patterns of connections across changes in group membership [1]. If changing conditions force individuals to “rewire” their connections, the social organization will change, but if individuals reconstruct similar social connections, the social organization will remain resilient. Therefore, an essential aspect of understanding animal social systems is describing how behavioral decisions allow for patterns of social organization to be continually reconstructed across changing contexts.

Social niche construction occurs when individuals take an active role in shaping their social environment [2]. When group conditions change, individuals face a new landscape of choices regarding who and how to interact. The decisions individuals make during these periods will simultaneously shape individual “success” in the group and collectively shape patterns of social organization that define the group. Although many factors outside social choices–such as resource distribution, differences in body size, differences in activity budgets, personality differences, and ecological factors [3]–play causal roles in shaping group formation and organization, investigating social niche construction requires isolating if social preferences and choices shape the emergence of group level patterns.

In changing group contexts, interactions can quickly shift from positive to neutral to harmful, depending on whom you interact with and how you interact with them. Among the most critical choices individuals make are the ones that minimize the risks of harmful interactions. Some individuals may be behaviorally incompatible due to differences in interaction styles, personalities, and motivations, or due to an increased risk of aggression resulting from specific interactions. Therefore, the ability to quickly assess behavioral compatibility and avoid incompatible conspecifics may be an essential aspect of navigating social interactions during fission-fusion changes [4], [5].

Differences in behavioral compatibility may cause individuals to selectively withdraw from or approach specific individuals within the group [6]. While stable approach-withdrawal preferences can minimize personal risks, it is unclear if they play a causal role in shaping the emergence of social organization. In some cases, morphological and physiological differences cause individuals to synchronize their behavior, resulting in social organization without strong social preferences [7]. For example, differences in age and sex often correspond to differences in size and morphology. In turn, these differences can lead to assortment as individuals converge on different habitats due to their varying metabolic and physiological needs [8], [9]. Thus, patterns of assortment can be passive, and occur when individuals lack any approach-withdrawal preferences for individuals in the group.

Nonetheless, patterns of assortment can also be active. Behavioral incompatibilities may cause individual to be so selective in their interactions that it shapes the overall social organization. Saltz et al. 2011 found that the social organization of *Drosophila melanogaster* groups was collectively shaped by the presence of aggressive males [10]. Populations with more aggressive males were characterized by higher rates of displacement, resulting in smaller, less cohesive groups, in contrast to populations with fewer aggressive males. Sapolsky (2004) documented that after aggressive males were removed from an olive baboon (*Papio anubis*) troop, females modified how they interacted with new males in a way that created a less assorted, more egalitarian organization in the troop [11].

In many contexts, some of the most pronounced behavioral incompatibilities exist between the sexes. In particular, studies have shown that aggressive interactions between males and females can result in substantial risks to females [12] even during non-reproductive periods. As such, females often employ behavioral strategies to control interactions with males to minimize harassment from them. Female-driven sex assortment has been documented in social mammals and increasingly in birds and invertebrates. In particular, females tend to have strong associations with other females, whereas males disperse to neighboring groups [13]. In some mammals [3], birds [5], [14], and insects [15], females form cohesive female subgroups within the larger group.

This study describes the social organization of captive flocks of Gouldian finches to identify areas where social niche construction may be occurring. Previously, it had been demonstrated that Gouldian finches exhibit age-based assortment patterns [16], with juveniles maintaining strong connections with siblings as they transition to independence. This study aims to determine whether patterns of social assortment are maintained across adulthood and to identify the behavioral mechanisms that sustain them. In particular, we build on previous findings in different bird species [4], [14] to investigate whether individuals exhibit consistent preferences for approaching and withdrawing from certain conspecifics across varying flock conditions.

Gouldian finches are an endangered estrildid finch that inhabits the subtropical grasslands of northern Australia. Little is known about the social organization of Gouldian finch flocks, but their fluidity has led some to speculate they lack within-group structure [17], [18]. In the wild, large fission-fusion flocks composed of hundreds of individuals are commonly observed. The sex ratio of Gouldian flocks is biased towards males [19], [20], and individuals in captivity and wild maintain family connections with siblings during early life [16], [21].

Gouldians do not exhibit sexual dimorphism. Males and females are similar in body size, shape, and structure [22], [23]. Across both sexes, Gouldians do exhibit a color polymorphism where adults show discrete differences in head coloration. Differences in head coloration correspond with differences in behavior and personality. Red-headed individuals are more aggressive, with fights between red-headed individuals being more intense than those involving other head colors, and red-headed females displacing black-headed females more often [24]. In the wild, black-headed individuals are the first to descend to waterholes [23]. In captivity, some studies have shown that black-headed individuals were more likely to approach and interact with a novel object faster and return to a food bowl sooner after the presentation of a simulated predator [25], [26], while others have found that red-headed individuals are more neophilic in other contexts [27].

It is currently unknown how different color morphs and sexes associate with each other in a semi-naturalistic context. Due to behavioral differences across the sexes and color-morphs, it is hypothesized that Gouldian finch flocks will show a significant assortment by sex and color-morph status. Research on animal social networks often focuses on associations, where social connections are measured when two or more individuals inhabit the same place in close proximity. While associations can be powerful measures of social organization, the social choices that build connections can be difficult to measure without observing directed interactions between known individuals. By recording interaction preferences, we can uncover how social choices give rise to higher-order patterns of organization [28]. This paper aims to describe the stable social preferences that persist across flocks differing in size and composition, with the goal of pin-pointing potential generative behavioral mechanisms underlying social network organization in Gouldian flocks.

## 2 Methods

### Subjects

All individuals in this study were Gouldian finches. Finches were acquired from professional breeders in Arizona and Florida, U.S.A., or bred in the lab. Each bird was provided with a uniquely colored leg band for individual recognition. Males, females, and juveniles were distinguished from each other based on plumage and behavioral differences, with adult females exhibiting more subdued chest coloration compared to males. A total of 62 individuals were used in this study. During 2020-2021, we used 32 birds, comprising 10 females and 22 males. During the 2022-2023 period, we used 30 birds, comprising 11 females and 19 males. Head color was readily observed in adults and assessed in juveniles at the time of their first molt after the end of the data collection period. Juveniles were distinguished from adults on the basis of their gray head, olive back, and lack of typical adult plumage. In this study, all juveniles were under one year of age since fledging the nest, and adults were older than a year and exhibited adult plumage.

All birds had a daily diet consisting of finch seed mix (Higgins finch seed mix) and soaked Perle Morbide pellets, with eggshell, boiled eggs, and dandelion and mustard greens provided twice a week. Fresh water was continually available.

### Aviaries

We used a series of two aviaries in this study. Each aviary consisted of identical dimensions (23 × 72 × 41 in) and similar environmental conditions. The environment within the aviaries was equivalent, with natural eucalyptus and plastic privacy perches, small shrubs, a feeding station, and a bowl for water baths.

### Data Collection

Focal sampling was used throughout this study. Each focal block lasted 5 minutes, and during each block, one individual was continually monitored, recording a list of behaviors that occurred within a 2-inch radius (approximately 1 body length). During focal sampling, we recorded approaches, leaves, displacements, and other behaviors (see the supplementary material). All observations were conducted using voice recognition technology, enabling the observer to maintain continuous visual contact with the focal bird throughout the sampling block. An approach was scored when another individual approached another within a 2-inch radius. Before data collection, a video of marked perches was made to establish that 2 inches was the typical social distance observed between two resting adult finches. GMK collected all focal data.

### Procedure

This study was carried out in two sessions. The first session, which occurred from 2020 to 2021, consisted of fission-fusion perturbations involving the sequential introduction of juveniles into the flock. The first session came from a study on juvenile social integration published elsewhere [16], but the data on overall social organization is analyzed here for the first time. The second session occurred in 2022 and investigated if stable patterns of social organization persist across a large-scale group perturbation in a larger adult flock.

#### Session One

Baseline: A flock of unfamiliar adults was placed together in a single aviary on August 5th, 2020, and allowed 4 days to acclimate. From August 7th, 2020 to August 17, 2020 focal samples were collected n the morning between 7:45 to 10:00 AM. A total of 352 focal samples were taken to measure approach behavior. Each individual was observed for 32 blocks, totaling 160 minutes each.

First Fission-Fusion: The flock from baseline was maintained, but on October 15, 2020, four novel juveniles (40 days post-fledging) were introduced into the aviary. From October 6, 2020 to December 12, 2020 focal samples were taken each morning between 7:00-10:00 AM. A total of 503 focal samples were collected, with each bird sampled across 34 blocks, except for one bird, which was observed for 26 blocks due to temporary removal from the aviary for illness.

Second Fission-Fusion: The flock from the previous fusion condition was split into two flocks on February 10, 2021. Aviary one contained three adult males, two adult females, and two juveniles, while aviary two contained four adult females, two adult males, and one juvenile. Immediately after splittin,g three juveniles were also introduced to each aviary. From February 11 to February 20, 2021, a total of 84 blocks were recorded between 7:30 and 10:30 AM. Each individual in the flock was recorded in eight blocks, totaling 40 minutes each.

Third Fission-Fusion: The flocks from the previous Fission-Fusion sections were maintained until May 25, 2021. On that date, a group of five novel juveniles was introduced into aviary 1. On June 11, 2021, the five juveniles were transferred to aviary 2. One juvenile had to be removed from Aviary 2 due to illness, and it was replaced with an adult female on June 10, 2021. Focal samples were collected from May 26 to June 27, 2021, between 7:00 and 9:45 AM. A total of 240 blocks were collected in each aviary, with each individual sampled for a total of 16 blocks each or 80 minutes of observations.

#### Session Two

Baseline: Birds from session one were removed and placed in the holding aviary, and a new group of birds was introduced to the aviaries. Both aviaries contained 6 females and 9 males. From January 31 to March 10, focal samples were conducted between 7:00 and 9:45 AM. A total of 510 for a total of 17 blocks per individual, for 85 minutes per individual.

### Male Swap

On June 7 2022, all males were swapped between the two aviaries. From June 8 to July 2, observational focal samples were collected. Each individual was sampled 17 times for 85 minutes of observation.

### Analysis

Social interactions are inherently nonindependent, so our analysis combined generative network models, permutation tests, and Bayesian multimembership models. We first analyzed the social network structure using permutation methods and community detection algorithms. Next, the individual social preferences underlying these group patterns were analyzed using Bayesian generalized linear mixed models. All analyses were conducted in R version 4.4.1.

### Modeling Social Network Structure

One aim of this paper is to investigate whether Gouldian finch flocks show group-level patterns of social assortment. To test this, we calculated a sex-based weighted assortativity coefficient first proposed by Leung and Chau [29]. To test the significance of these coefficients, we used permutation-based methods. Sex was randomized for all individuals in the flocks while the edge structure of the network was preserved. We conducted 10,000 permutations on each network, resulting in a null distribution of assortativity coefficients for each flock. Empirical p-values were then calculated by looking at the proportion of assortativity coefficients from the null distribution that were larger and smaller than the observed assortativity coefficient for that network.

Further modeling of the social network’s structure was conducted using spinglass clustering algorithms. This algorithm identifies a subgroup within the larger network by optimizing a function that rewards connections within the subgroup and penalizes those between subgroups. This algorithm was chosen because it allows for the assessment of subgroup structure in small networks with weighted and directed ties [30].

### Modeling Social Preferences

To model individual social preferences, we used a Bayesian generalized linear mixed model with a multi-membership term using the Bayesian program STAN in the package *brms* [31]. These models are particularly suitable for analyzing social preferences within freely interacting groups, as they allow for random-effect structures that address the types of dependencies inherent in relational social data and repeated sampling of individuals. These models have previously been applied to the study of social relationships in animal groups with small sample sizes [32]. All models were run for 10,000 iterations across two chains with a warm-up period of 2,000 iterations. The significance of fixed effects was evaluated using the 95% credible intervals. If the credible interval did not overlap zero, the effect was considered significant. To ensure model convergence, we inspected trace plots and applied Heidelberger and Welch diagnostics to verify that the Markov chain reached a stationary distribution.

We conducted two main models: one that examined the distribution of approaches and the other that analyzed the distribution of leaves. The dependent variable was either the frequency of approaches or leaves between each dyad in the flock. Each dyad was categorized according to whether the initiator and recipient shared the same sex, family, or head color. These categories were incorporated as fixed effects in the model. Random effects accounted for variations between aviaries, the introduction date of the initiator to control for introduction order, and a multi-membership term that included both the identity of the initiator and receiver to address the non-independence of dyads. To evaluate the direction of average approach rates (calculated as the number of approaches directed toward individuals divided by the number of individuals), we employed permutation-based Wilcoxon signed rank tests, adjusting for multiple comparisons using the false discovery rate.

A third model was restricted to the 2022 male swap study. This model was identical to the ones described above, but had an additional explanatory variable containing whether the data came from before or after the male swap between the aviaries.

### Sub-Group Stability

To investigate if individuals maintained the same associations within same-sex subgroups, we correlated the weight of each dyad before and after the male swap in the 2022 dataset. To control for the non-independence of dyads, we used a permutation test for correlation coefficients in the package *PermCor*. If individuals interacted with the same individuals in the same fashion, a positive correlation should be shown. Empirical p-values for correlation coefficients were calculated for both male and female sub-groups by comparing the observed correlation coefficient to the permuted null distribution.

**Figure 1:**
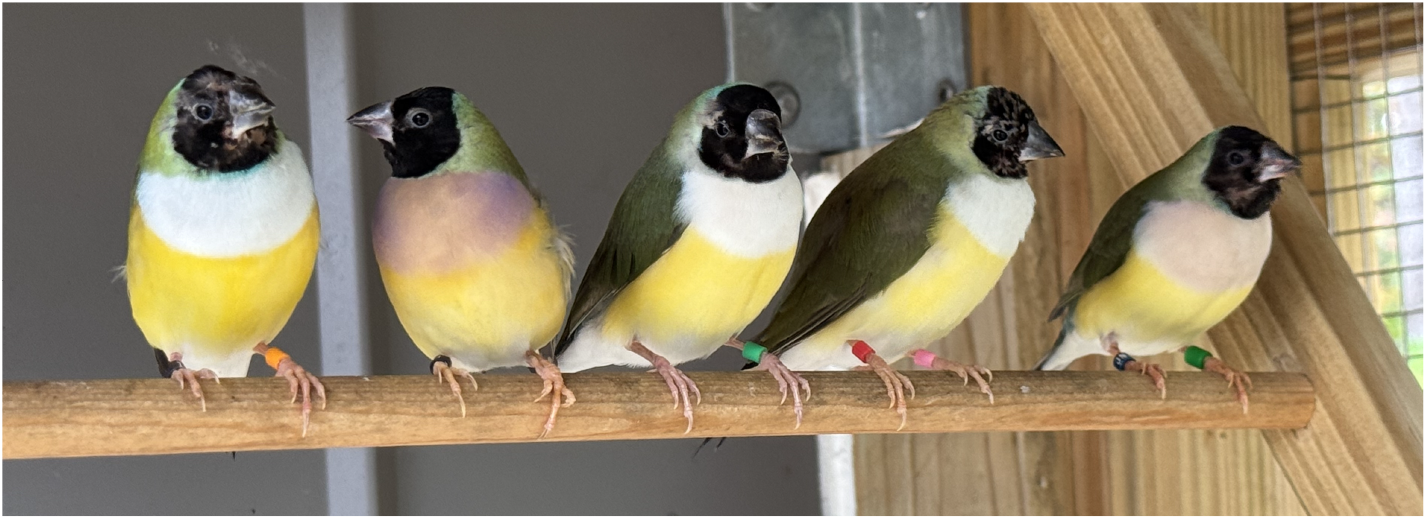
Picture of temporary female subgroup

## 3 Results

### Social Approaches and Leaves

A total of 4,916 approaches were recorded during the baseline period, including 236 leaves. During the first fission-fusion period, a total of 8,874 approaches and 355 leaves were recorded. During the second fission-fusion period, we recorded a total of 2184 approaches and 94 leaves were observed in aviary 1, and 2345 approaches and 106 leaves were observed in aviary 2. In the third fission-fusion period, we recorded a total of 4425 approaches and 180 leaves in aviary 1, and 3991 approaches and 180 leaves in aviary 2. In the second session, we recorded a total of 2883 approaches and 123 leaves in Aviary 1, and 3330 approaches and 143 leaves in Aviary 2 during baseline, and in the male swap period, we recorded a total of 1947 approaches and 39 leaves in Aviary 1 and 2464 approaches and 80 leaves in Aviary 2. We found no significant differences in the number of approaches or leaves between males and females (Supplementary Materials).

### Social Network Structure

Across all periods, our spin-glass clustering algorithms identified separate subgroups within the larger network. In all but one flock (Aviary 1, Second Fusion 2021), one cluster emerged that contained mostly adult females (Table 1). Weighted assortment coefficient values were above zero for all flocks, indicating that networks were characterized by more connections within sex classes than outside them (Table 1). Permutation tests revealed that, in most flocks, these sex-based assortment coefficients were significantly higher than the permuted null distribution (Table 1).

**Table 1:** Spin-glass Communities and Assortment Coefficients. The first two subgroups identified by the algorithm are listed in column subgroup 1 & 2. F is adult female, M is adult male, JM is juvenile Male, and JF is juvenile female. The assortment coefficient is calculated by sex, with values above 0 indicating positive assortment. P-value is the permutation-based empirical p-value based on 10,000 network permutations.

| Aviary | Session | Subgroup 1 | Subgroup 2 | M-F Assort | p-value |
| --- | --- | --- | --- | --- | --- |
| Aviary 1 | Baseline 2020 | 6 F | 5 M | 0.23 | 0.04* |
| Aviary 1 | Fusion 2021 | 6 F | 5 M | 0.24 | 0.04* |
| Aviary 1 | Fission-Fusion 2021 | 2F & 1M | 3 JM | 0.24 | 0.03* |
| Aviary 2 | Fission-Fusion 2021 | 4 F & 1M | 4 JM | 0.31 | 0.06 |
| Aviary 1 | Second Fusion 2021 | 3 F & 1 JF & 2 M | 4 JM | 0.17 | 0.01* |
| Aviary 2 | Second Fusion 2021 | 4 F & 2 M | 9 J | 0.43 | 0.03* |
| Aviary 1 | Baseline 2022 | 7 M | 5 F | 0.24 | 0.02* |
| Aviary 2 | Baseline 2022 | 8 M | 7 F & 1M | 0.28 | 0.03* |
| Aviary 1 | Swap 2022 | 5 F | 5 M | 0.27 | 0.02* |
| Aviary 2 | Swap 2022 | 6 F | 5 M | 0.26 | 0.03* |

### Social Preferences

Our model for approaches revealed a significant influence of sex and age on the frequency of interactions between each dyad (Table 3). Post hoc analysis showed that females and males did not differ in their overall approach rates (Wilcoxon rank sum exact test: Average Male Approach Rate = 17.85957, Average Female Approach Rate = 21.67537, W = 308, p-value = 0.1181). When looking at the approach rates directed towards females, males had significantly lower approach rates towards females than other females (Wilcoxon Rank Sum Test: Mean Female-Female = 32.17, Male-Female = 18.86, W = 637, p = 0.0004345), and males had higher approach rates towards males than females (Wilcoxon Rank Sum Test: Mean Male-Male = 16.85, Mean Female-Male = 11.18, W = 242, p = 0.009).

**Table 3:**
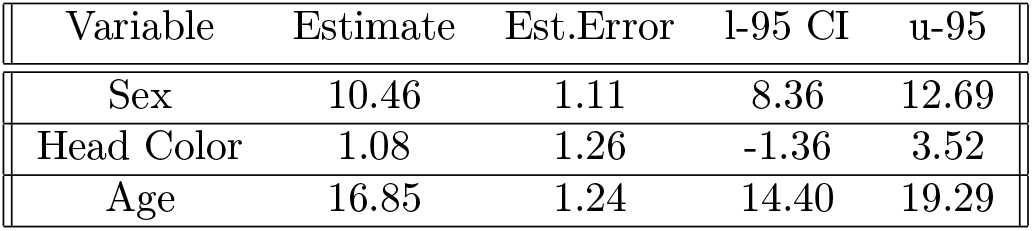
Results of the Bayesian regression model on approaches.

Within each sex, females had higher approach rates directed towards females in comparison to approach rates towards males (Wilcoxon signed rank test: Mean Female-Female = 32.17, Mean Female-Male = 11.18, V = 228, p < 0.0001), but in males we found no differences in approach rates directed towards males or females (Wilcoxon signed rank test: Mean Male-Female = 18.8616, Mean Male-Male = 16.85754, V = 45, p = 0.41).

**Figure 2:**
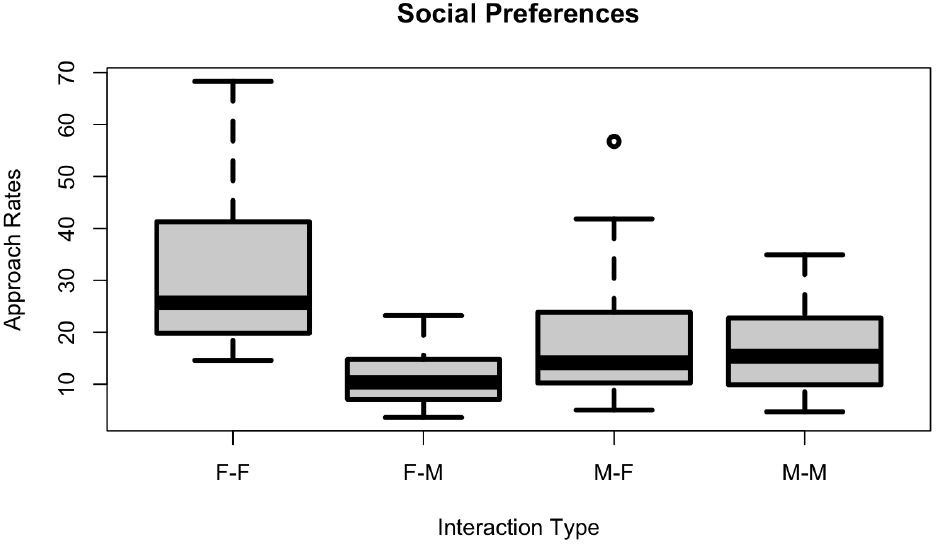
Sex-based Preferences

Our second model examined the factors that influenced the frequency of an individual’s leaving after being approached. Our model for leaves showed a significant influence on sex, head color, and age, with 95-percent credible intervals for each not overlapping with zero (Table 4). The negative credible intervals for sex and head color did not overlap with zero, suggesting that leaves were more likely to occur between different-sex or different head color dyads than same-sex dyads.

**Table 4:**
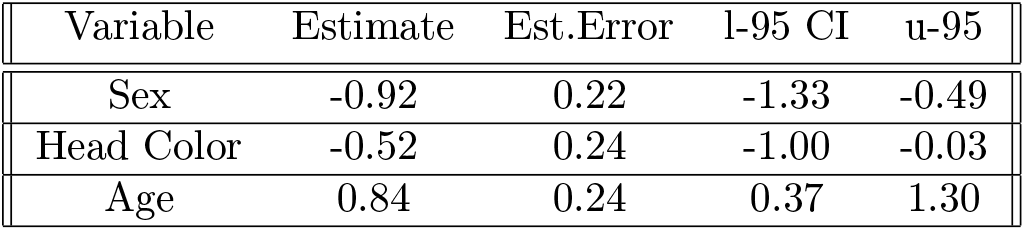
Results of the Bayesian regression model on Leaves.

Further analysis revealed that female social choices were the primary driver of this effect. Male approaches were more likely to be followed by leaves in contrast to female approaches (Wilcoxon Rank Sum Test: Female Approach-Leave = 0.311, Male Approach-Leave = 1.00, W = 684.5, p < 0.0001). There were no significant differences in leaves between males and females after being approached by a female (Fig. 3, Wilcoxon signed rank test: Female Approach-Female Leave Mean = 0.28, Female Approach-Male Leave Mean = 0.34, V = 78, p-value = 0.20). However, females were more likely to leave than males after being approached by a male (Fig. 3, Wilcoxon signed rank test: Male Approach-Female Leave Mean = 1.47, Male Approach-Male Leave Mean = 0.55, V = 695, p < 0.0001).

**Figure 3:**
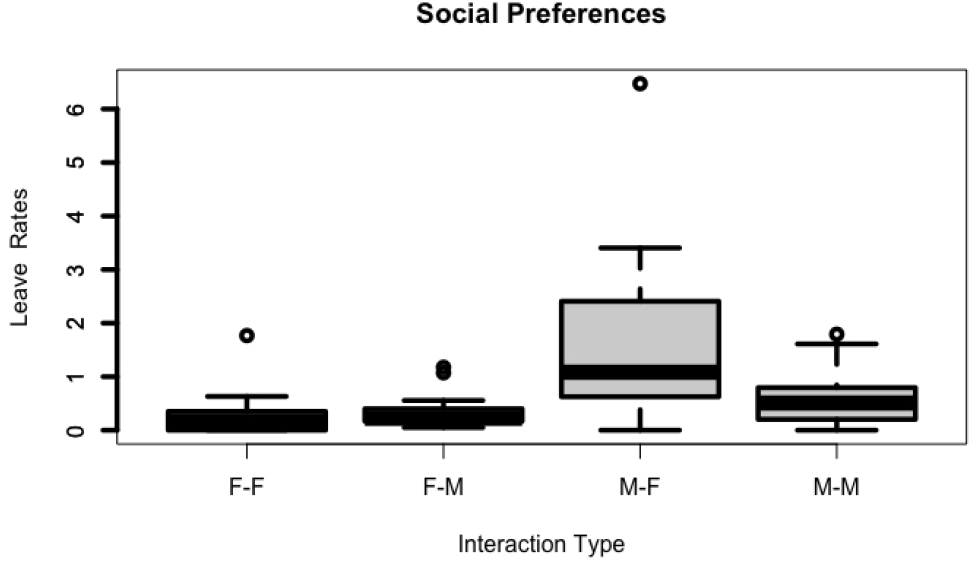
F-F Looks at rates of females leaving after female approaches-Female Leave, F-M is the rate of males leaving female approaches-Male Leave, M-F is the rate of Male approaches-Female leaves, and M-M is the rate male approaches-male leaves

### Models for 2022 Period

When we restricted our analysis to the 2022 period, we found equivalent results. Since all individuals in this session were adults, age was removed as an explanatory factor, and session (before male swap vs. after male swap) was added as an explanatory factor. Our model for approaches showed a significant influence of sex (Estimate = 0.55, Standard Error = 0.02, 95% CI = 0.51-0.59) and session (Estimate = −0.31, Standard Error = 0.02, 95% CI = −0.35 − −0.27) on the strength of dyads.

Overall interaction rates were lower after the males were swapped then before (Wilcoxon Signed Rank Test: Mean Interaction Rate Before = 14.8, Mean Interaction Rate After = 10.50, V = 359.5, p = 0.009). This was largely driven by a reduction in female approaches. Females showed lower interaction rates after males were moved across the aviaries (Wilcoxon Signed Rank Test: Mean Female Interaction Rate Before = 18.86, Mean Female Interaction Rate After = 11.50, V = 359.50, p = 0.02), whereas no significant change occurred in male interactions.

The rate of female-to-female approaches was significantly higher than female-to-male approaches both before (Wilcoxon signed rank test: Mean Female-female = 29.45, Mean Female-Male = 8.26, V= 66, p < 0.0001) and after male swaps (Wilcoxon signed rank test: Mean Female-female = 17.73, Mean Female-Male = 5.27, V= 67, p < 0.0001). Males did not show any significant differences before or after the swap.

Our model for leaves during the 2022 period did not show a significant influence of any variables. This could be due to the relatively smaller number of leaves that occurred during this period. However, Wilcox tests showed that females were significantly more likely to leave after being approached by males than by females both before (Wilcoxon signed rank test: Male Approach-Female Leave Mean = 1.7, Male Approach-Male Leave Mean = 0.34, V = 148.5, p < 0.0007) and after (Wilcoxon signed rank test: Male Approach-Female Leave Mean = 1.5, Male Approach-Male Leave Mean = 0.27, V = 115.5, p = 0.015) the male swap period.

### Sub-Group Stability

Across the baseline and male swap period we found significant correlations between the strength of dyads within female sub-groups (rho = 0.61, p = 0.01) and within male sub-groups (rho = 0.33, p < 0.001).

## 4 Discussion

This study described the fine-scale structure of social organization in Gouldian finch flocks. Our results showed that sex-based assortment is a robust feature of Gouldian social structure. Sex assortment was sustained across flocks that differed in age, sex, stability, familiarity, and size. Furthermore, these patterns were maintained across changes in group size and composition, with significant levels of sex assortment occurring in nearly every flock we observed. While most flocks contained both male and female-specific subgroups, we found that females maintained higher levels of female-to-female approaches than expected. In contrast, males did not show a significant preference for either males or females. Furthermore, females were significantly more likely to immediately leave after being approached by a male, strongly suggesting that they prefer to avoid interactions with males over interactions with females.

The ability to navigate the risks and benefits of interactions is essential for success in the group [13]. There can be significant risks when interacting with novel or aggressive conspecifics, and individuals may buffer against these interactions by becoming selective in whom they approach [1]. Some interactions can have significant benefits, such as access to beneficial social information or experiences necessary for the development of competent behavioral abilities [33] (West & King, 2002). In such cases, individuals may be selective and preferentially interact with individuals due to the beneficial consequences of those interactions [34], [35]. To understand how individuals navigate the potential risks and benefits of interactions, we must first have a detailed description of how group members allocate their social decisions in the group to see which individuals are preferred or avoided.

To our knowledge, this study provides the first evidence for sex based-assortment within estrildid finches. Previous findings have shown that juvenile Gouldian finches form subgroups with their siblings in both captivity [16] and the wild [21], suggesting that patterns of social organization found in captive contexts may generalize to the wild. While sex assortment is increasingly being discovered in birds, this study is among the few that show that sex assortment persists across dynamic group changes. In Brown-headed Cowbirds (*Molothrus ater*) females preferentially associate with other familiar females across significant scale changes in group size and composition [4]. Together, these findings suggest that females may be a strong driver of social organization in the fission-fusion societies that characterize many bird species.

Despite sex assortment and segregation being increasingly observed in a wide range of animal species [3], the exact processes by which individuals come to surround themselves with same-sex individuals are often unknown. In some species, this process seems to be entirely passive, with behavioral decisions and preferences playing no role in driving the formation of separate groups or communities. Specifically, passive assortment has been proposed as a significant cause of sexual segregation in ungulates, where differences in metabolic needs, size, or activity patterns create self-organized sex-based groups outside individual social preferences. For example, in feral goats (*Capra hircus*), higher temperatures will cause males to end resting earlier to begin foraging [8]. This results in increased synchrony among males and dis-synchrony between males and females, leading to complete segregation. Similar patterns are also seen in sexually dimorphic bird species. In arctic buntings, males tend to occupy colder areas as their larger size allows them to regulate their temperature better when faced with severe winter weather [9].

Although social assortment may be influenced by a wide range of factors, our results strongly suggest that sex assortment is an active process in Gouldian Finches. By ‘active,’ we mean that the social approach-withdrawal preferences play a causal role in the formation of sub-group structure. Both male and female Gouldian finches are similarly sized, making it less likely that differences in activity synchrony or metabolic needs drive assortment [22], [23]. Captive settings also minimize assortment based on habitat choice or differences in metabolic demands, leaving patterns of social association more likely to be the result of social decisions.

In many cases of female-driven assortment, a common factor is the avoidance of males. Male interactions can carry increased risks of aggression and harassment in many species, and individuals may act to avoid these by forming female-only groupings. Our finding that females showed an increased propensity to leave after being approached by males strongly suggests that females actively avoid male interactions. These findings align with findings in other species. For instance, in guppies, patterns of male harassment have been shown to drive females into female-only subgroups [15]. Similarly, in cockroaches (*Diploptera punctata*), females cluster together and actively displace males from their subgroups. In Siberian Ibex *Capra sibirica* sexual segregation increased as males ages and their more aggressive behavior became incompatible with females [36]. In pheasants *Phasianus colchicus*, after hatching, males and females showed no differences in size or activity budgets, but females still prefer to engage with other females [14]. Together, these studies suggest that differences in behavioral compatibility shape preferences for interacting with members of the opposite sex. Furthermore, these preferences in turn shape the structure of the grou,p resulting in stable patterns of assortment within an otherwise cohesive flock.

By avoiding potentially disruptive male interactions, females may construct a more favorable proximal social environment for themselves. Our flocks were biased towards males to match the sex ratio found in many flocks in the wild and captivity [19]. In such settings, close relationships among females may create a social buffer that leads to lower stress levels and better fitness outcomes. Studies in primates and birds have shown that strong connections between individual females may have substantial fitness benefits. In baboons, stronger female bonds are associated with higher offspring survival and increased female longevity [37]. In Brown-headed cowbirds, females with stronger preferences for familiar individuals during the fall were more likely to lay more eggs during the subsequent breeding season [5]. Our findings suggest that female Gouldians were able to maintain connections with specific females across changes in group composition. This opens the possibility that within female sub-groups, females sustain enduring relationships with other females. Subsequent work will investigate whether the characteristics of these relationships may shape later reproductive success and stress responses.

Females may also associate with other females to access valuable social information. Cues displayed by females can communicate mate and song preferences and exposure to these cues can shape reproductive, competitive, and social decisions in females [38]–[40]. In brown-headed cowbirds, females acquire mate preferences through interactions with other females, and females who were more familiar with each other during the fall are more competitive with each other during the breeding season (Kohn, in prep). Cowbird flocks also segregate according to age and sex in response to specific stimuli, such as playback of specific songs [41]. While little is known about the mate preferences of Gouldian finches, relationships between females may allow females to monitor the preferences of others. In one of the flocks, two of the strongest connected females (FYER and FB2G) were both paired with the same male (MBNB). Furthermore, male-to-female aggression was rarely observed. Across all flocks, only 44 incidents of fights were observed, and these did not significantly differ between males and females (Kohn, personal communication). While further research should investigate the broader consequences of female-to-female interactions, our findings suggest that female subgroups could be used to access information from other females.

Unlike females, males did not show any significant social preferences. Nonetheless, our social networks show clear evidence of male communities via our spinglass clustering algorithm, especially during the 2022 data collection period. Together, these results suggest that male groups were primarily caused via exclusion by females. Nonetheless, within these male subgroups, the number of dyads was correlated across the 2022 groups. This suggests that males were able to maintain individual social relationships with other males despite changes in group composition. Further studies will investigate the differences in the emergence and function of individual relationships between males and females.

Social niche construction occurs when individuals actively shape the social environments they inhabit. Taken together, our results strongly suggest female Gouldian finches engage in social niche construction by creating a group-level pattern of sex assortment. Furthermore, our results show that female social decisions contribute to the robustness of this social structure, enabling it to persist across the changing group dynamics typical of Gouldians and many other species. These results contribute to the growing list of examples where social niche construction enables both the creation of new social systems and the persistence of existing social systems despite changes in overall group dynamics and ecological shifts. The implications of this are vast, as they suggest that social factors can become the primary factor shaping the selective environment individuals experience, and highlight the importance of studying female interactions as a distributed system shaping the organization of fission-fusion societies.

## 5 Acknowledgments

This study was supported with a grant from the Van Vleck Award. All research was done in accordance with ASAB/ABS Guidelines for the Use of Animals in Research and was approved by the University of North Florida’s Institutional Animal Care and Use Committee protocol number 20-006.

## Notes

### Competing Interest Statement

The authors have declared no competing interest.

